# Determining Best Practice for the Spatial Poisson Process in Species Distribution Modelling

**DOI:** 10.1101/2023.01.10.523499

**Authors:** Sean Bellew, Ian Flint, Yan Wang

## Abstract

Poisson processes have become a prominent tool in species distribution modelling when analysing citizen science data based on presence records. This study examines four distinct statistical approaches, each of which utilises a different approximation to fit a Poisson point process. These include two Poisson regressions with either uniform weights or the more elaborate Berman-Turner device, as well as two logistic regressions, namely the infinitely weighted logistic regression method and Baddeley’s logistic regression developed in the context of spatial Gibbs processes. This last method has not been considered in depth in the context of Poisson point processes in the previous literature. A comprehensive comparison has been conducted on the performance of these four approaches using both simulated and actual presence data sets. When the number of dummy points is sufficiently large, all approaches converge, with the Berman-Turner device demonstrating the most consistent performance. A Poisson process model was developed to accurately predict the distribution of Arctotheca calendula, an invasive weed in Australia that does not appear to have been the subject of any species niche modelling analysis in the existing literature. Our findings are valuable for ecologists and other non-statistical experts who wish to implement the best practices for predicting species’ distribution using Poisson point processes.

## 1 Introduction

Species Distribution Modelling (SDM) concerns the use of environmental covariates to model the presence of a species over a specified domain, predicting the geographical distribution of a species over time (Hof et al., 2012). This outcome is typically achieved using information from a species presence and pairing this with known environmental covariates at those locations (Guillera-Arroita et al., 2015). SDMs help identify relevant environmental factors which affect species’ distribution, allowing a better understanding of how different events influence the species, as well as providing insight into how it may be affected under future environmental conditions, providing value in the fields of ecology, biogeography, biodiversity conservation, and natural resource management (Guillera-Arroita et al., 2015).

Presence-only (PO) data sets contain only sites where a species has been recorded as present; there is no additional data on whether the species is present or absent at other sites in the study area. These data sets are opportunistically sampled and are not obtained through structured studies that produce presence-absence (PA) or occupancy-detection data sets. Such data sets are commonly found in atlases, museums, herbariums and incidental observation such as citizen science datasets (Pearce and Boyce, 2006), and have been used in 53% of SDM publications (Guillera-Arroita et al., 2015). PO data is sometimes accompanied by a background sample. The term background is in reference to locations where the presence or absence of the species has not been recorded, but that are included to convey additional information about the environmental features of other locations to the model.

PO data consists in point events – a set of point locations where a species has been observed. The example data used in this paper comprises 832 point locations of *Arctotheca calendula*, commonly known as Capeweed. This species occurs in Southern Australia. It is a pest and competitor to legume crops and leads to weight reduction for livestock in pastures where it is present (Brundu et al., 2015; Conning et al., 2011). However, Capeweed does not appear to have been the subject of any SDM analysis in the current literature. In our study the presence-only data used spanned 2010-2021 and focused on Southern Australia (GBIF, 2021). We included various environmental variables to identify the key environmental factors driving the distribution of *Arctotheca calendula*.

In species distribution modelling, the Poisson process model (PPM), has been proposed as an natural way for analysing presence-only data (Chakraborty et al., 2011; Renner and Warton, 2013; Warton and Shepherd, 2010). PPMs have been shown closely related to other widespread methods in ecology, such as MAXENT (Aarts et al., 2012; Fithian and Hastie, 2013; Renner and Warton, 2013; Wang and Stone, 2019), logistic regressions (Baddeley et al., 2010; Warton and Shepherd, 2010) and other statistical methods (Aarts et al., 2012; McDonald et al., 2013; Wang and Stone, 2019). Different methodologies and relevant approaches are available to fit the Poisson PPM, including the conventional likelihood approach (Warton and Shepherd, 2010) and infinitely-weighted logistic regression (IWLR) (Fithian and Hastie, 2013). Different quadrature schemes for approximating the integral in the likelihood function have been devised. The default in the `spatstat` package is the Berman-Turner device (Berman and Turner, 1992) (see Section 2 for details). Renner et al. (2015) thoroughly reviewed point process models, some of their advantages and some common methods of fitting them to presence-only data.

Baddeley et al. (2014) proposed a conditional logistic regression to fit spatial point processes, referred to as Baddeley’s logistic regression (BLR) in this paper. Instead of using a dense grid of pixels or dummy points, they generate a smaller number of dummy points at random locations. Given only the data at these (presence plus dummy) locations, the log-linear Poisson point process model is approximated by a logistic regression model, which can then be fitted using standard software. It has been asserted that this methodology performs better than the Berman-Turner device for a smaller number of dummy points, which would, as a result, make it more efficient (Baddeley et al., 2014). In this manuscript, we are interested in the performance of the BLR in comparison to the other methods that are typically employed in PPM fitting. We have not come across any prior research on this topic.

In this paper, the effectiveness of several distinct methods to fit the Poisson point process is evaluated and compared. The amount of dummy points required when studying PO data as well as the rate at which these methods converge are the primary questions of interest for our study. There were a total of four different models/approaches that were investigated, namely the gridded Poisson PPM, the weighted Poisson PPM with Berman-Turner device, the infinitely-weighted logistic regression (Fithian and Hastie, 2013) and Baddeley’s logistic regression. The paper led to the discovery of some new information and some recommendations. These recommendations help inform the choice of methodology used to fit SDMs with PO data. Our goal is to improve usability, which has been noted as a weakness in the SDM field (Illian and Burslem, 2017).

## 2 Methods

### 2.1 Poisson Point Process Model

Warton and Shepherd (2010) first proposed the use of Poisson point processes in SDM. A Poisson point process Møller and Waagepetersen (2004) *N* with intensity λ(*s*) on a study area *S* (*s* ∈ *S*) is characterised by the fact that the number of points in any measurable subset *B* ⊂ *S* is Poisson distributed *N*(*B*) ~ Po(λ(*B*)) with rate λ(*B*):= *∫_B_* λ(*ds*). As one of its fundamental properties, the Poisson point process’ number of points in two disjoint areas is independent. In SDMs, the intensity function λ(*s*) of the Poisson point process is typically modelled as a log-linear function log(λ_*β*_(*s*)) = *β’X*(*s*) for a given vector of covariates *X* computed at a location *s* and regression parameters *β* (Renner et al., 2015). Given observed locations {*s_1_*,…, *s_n_*}, the log-likelihood is given by (Renner et al., 2015)

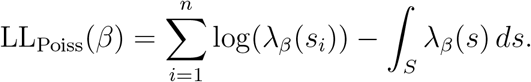

The integral appearing in this log-likelihood in practice must be approximated; this is achieved through numerical quadrature with weights *W_j_* (Warton and Shepherd, 2010) such that

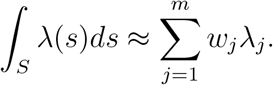

Warton and Shepherd (2010) noted that the quadrature points used in the integral approximation of the Poisson PPM likelihood are analogous to background points in a Poisson regression, and both methodologies give identical estimates as the number of quadrature points increases. It has also been shown that estimates derived by the Maxent method (Phillips et al., 2004), a widely-used machine learning based SDM method, is equivalent to point process estimates (Aarts et al., 2012; Fithian and Hastie, 2013).

### 2.2 Poisson regressions

There are several ways that one can approximate the integral appearing in the PPM log-likelihood (2.1) that take the form of a standard Poisson GLM regression. Here, we will study two different approaches when constructing the quadrature to fit the PPM likelihood function, namely, the gridded quadrature (PPM-GR) and the Berman-Turner device (PPM-BT). Both PPM-GR and PPM-BT maximize the likelihood function of the PPM, but they differ in their way of constructing the quadrature points. PPM-GR uses the simplest form where the study area is divided into *m* points distributed on a regular grid, and an equal weight of *w* = |*S*|/*m* is assigned to each quadrature point.

Berman and Turner (1992) generated the quadrature points by augmenting the presence points with a some dummy points. The quadrature weight *w_j_* associated with a presence or dummy point is the area of the corresponding tile obtained by a Dirichlet or Vorono tesselation. In this case, the PPM log-likelihood is approximated by

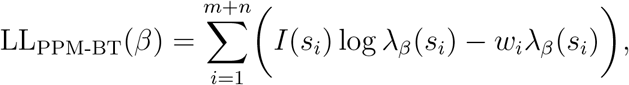

where *n* and *m* denote the number of presence and dummy points respectively, and *I* (*s_i_*) is the indicator function equal to 1 if *s_i_* is a presence point and 0 otherwise. This log-likelihood is equal to a weighted Poisson likelihood (Warton and Shepherd, 2010), and hence can be evaluated using GLM software. The method of PPM likelihood approximation with the Berman-Turner device (PPM-BT) is available to users through the `ppm` function available in the popular `spatstat` package (Baddeley et al., 2015) in `R`.

### 2.3 Logistic regressions

Different versions of a logistic regression have been proposed to fit a PPM. Given an ad hoc set of dummy points, the infinitely-weighted logistic regression (IWLR) (Fithian and Hastie, 2013) is set up via its log-likelihood given by

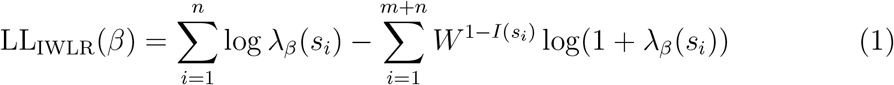

where *W* is a sufficiently large number. This method was shown to converge faster to the true parameter values compared to a standard logistic regression, and in particular performed better under a miss-specification of the model. Since the IWLR estimator is the solution of a weighted logistic regression, it can be implemented using standard GLM packages with weights manually specified (Renner et al., 2015).

Baddeley et al. (2014) proposed an alternative strategy to fit point process models using the conditional logistic regression. To avoid confusion with the standard logistic regression this method will subsequently be referred to as Baddeley’s logistic regression (BLR). The method relies on dummy points which, combined with the presence points, are fit according to a logistic regression model. The dummy points in this method are generated as a dummy point process with known intensity *ρ*, independent of the presence points, and distributed as a Poisson, binomial or stratified binomial point process (Baddeley et al., 2014). The proposed logistic likelihood is equal to

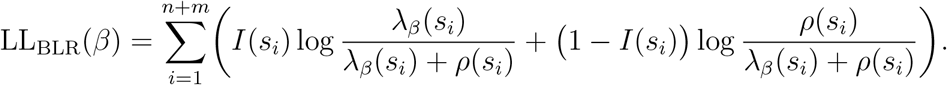

Baddeley et al. (2014) have shown that on average, *β* maximising LL_BLR_(*β*) also maximises the log-likelihood of the Poisson point process (2.1). Since LL_BLR_(*β*) is a standard logistic regression with offset term — log(*ρ*(*s_i_*)), it is readily fitted with GLM software (Baddeley et al., 2014).

## 3 Simulation Studies

A simulation study was carried out to compare the performance of the different methodologies presented in Section 2, namely, the infinitely weighted logistic regression (IWLR), Baddeley’s logistic regression (BLR), and two different schemes of quadrature points to fit the PPM likelihood function (PPM-GR and PPM-BT). In our study the virtual species’ distribution was simulated over a rectangular window within South-Western Australia, covering the area from 116° to 119° longitude and −34° to −31° latitude. Five bioclimatic covariates were assumed to drive the intensity function λ(*s*) of the underlying Poisson PPM for the species. These five covariates were selected out of 19 bioclimatic variables which have previously been used to fit plant species distribution models in this area and have been among the most commonly chosen covariates (Yates et al., 2010; Dalmaris et al., 2015; Thuiller, 2013). These five covariates are isothermality (BIO3), mean temperature of wettest quarter (BIO8), mean temperature of driest quarter (BIO9), precipitation of wettest month (BIO13), and precipitation of driest month (BIO14). They were selected to incorporate a variety of temperature and precipitation data and also to minimise the collinearity between covariates (Naimi et al., 2014). These covariates were sourced from the WorldClim database (Fick and Hijmans, 2017) and normalised to ensure the magnitudes of each were on the same scale. We used

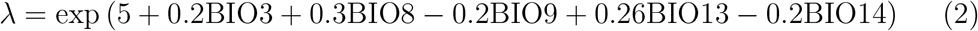

as the intensity function, with the parameters chosen so that the average number of points generated over the region is approximately 1000.

Poisson point patterns were simulated using the acceptance-rejection method (Pasupathy, 2011). To ensure a fair comparison between methodologies, identical sets of points were used to construct the dummy points for the PPM likelihood approximation as well as for the IWLR and BLR methods. These sets of dummy points were distributed on a uniform grid, which is the most straightforward choice for numerical quadrature.

The number of dummy points required to study PO data and how fast these different estimation methods converge are the primary questions of interest for our study. The number of dummy points in the fitting process varied from 49 to 8, 100 in our study. In practice, this was achieved by changing the grid step size, i.e., the distance between the two successive points on the grid, from the finest resolution with 8100 points to the coarsest with 49 dummy points. For each set of dummy points, 100 different realisations of the Poisson point process were generated and fitted with the four different methodologies. We then computed the root mean square error (RMSE) as

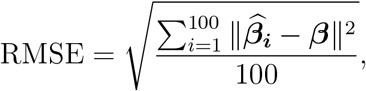

where ***β*** = (5, 0.2, 0.3, −0.2, 0.26, −0.2) are the true parameters in (2), 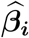 is the simulation-specific estimate, and || · || is the euclidean distance. In addition to RMSE, sample statistics of the coefficient estimates were computed and presented, in addition to the maximised log-likelihood value for each methodology. All simulations were carried out in R version 3.6.3 R Core Team (2021).

## 4 Results

Each of the parameter estimates, corresponding to the slope and intercept coefficients, was computed for each methodology for various numbers of dummy points (see Figure 1). Similarly, the maximised log-likelihood value and RMSE were recorded and are presented in Figure 2. We notice from the simulations that the estimates of the infinitely-weighted logistic regression (IWLR) proposed by Fithian and Hastie (2013) are nearly identical to the maximum likelihood estimates of the gridded Poisson PPM (PPM-GR), when dummy points in both methods were chosen on a uniform grid. The results show that the parameter estimates of all four methodologies converged to the true parameter values as the number of dummy points increases. However, they converge at different rates and in fact perform differently when the number of dummy points is small. In this circumstance, PPT-BT outperforms the three other methods. In the plot of the log-likelihood values in Figure 2, it appears that compared to the other methodologies, the log-likelihood for the PPM-BT does not vary much when the number of quadrature points increases. Furthermore, we see that the PPM-GR, PPM-BT, and IWLR all converge to the same value as the resolution increases. The maximised log-likelihood for the BLR, however, converged to a different value, and this likelihood has thus been linearly transformed on the plot for ease of comparison between the methodologies. The maximised log-likelihoods for the PPM-GR and IWLR methods depart slightly from the true value when the number of quadrature points is small. Using the RMSE as the measure for performance, as well as observation of the parameter estimates, we see that for small amounts of dummy points (roughly less than 400) the model fit for PPM-BT slightly outperforms BLR, and more significantly outperforms PPM-GR and IWLR.

**Figure 1:**
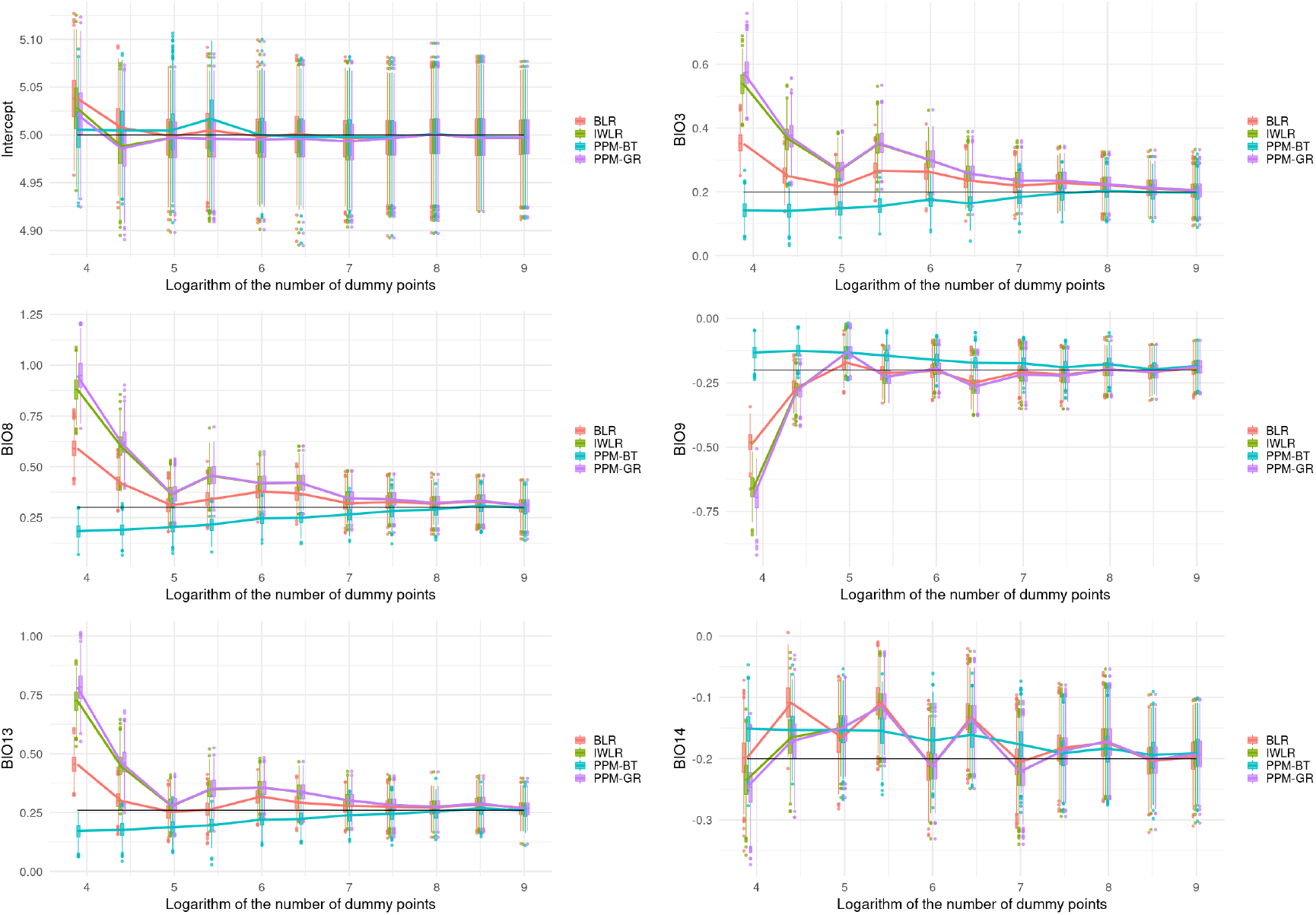
Parameter estimations of each regression coefficient for various numbers of dummy points.

**Figure 2:**
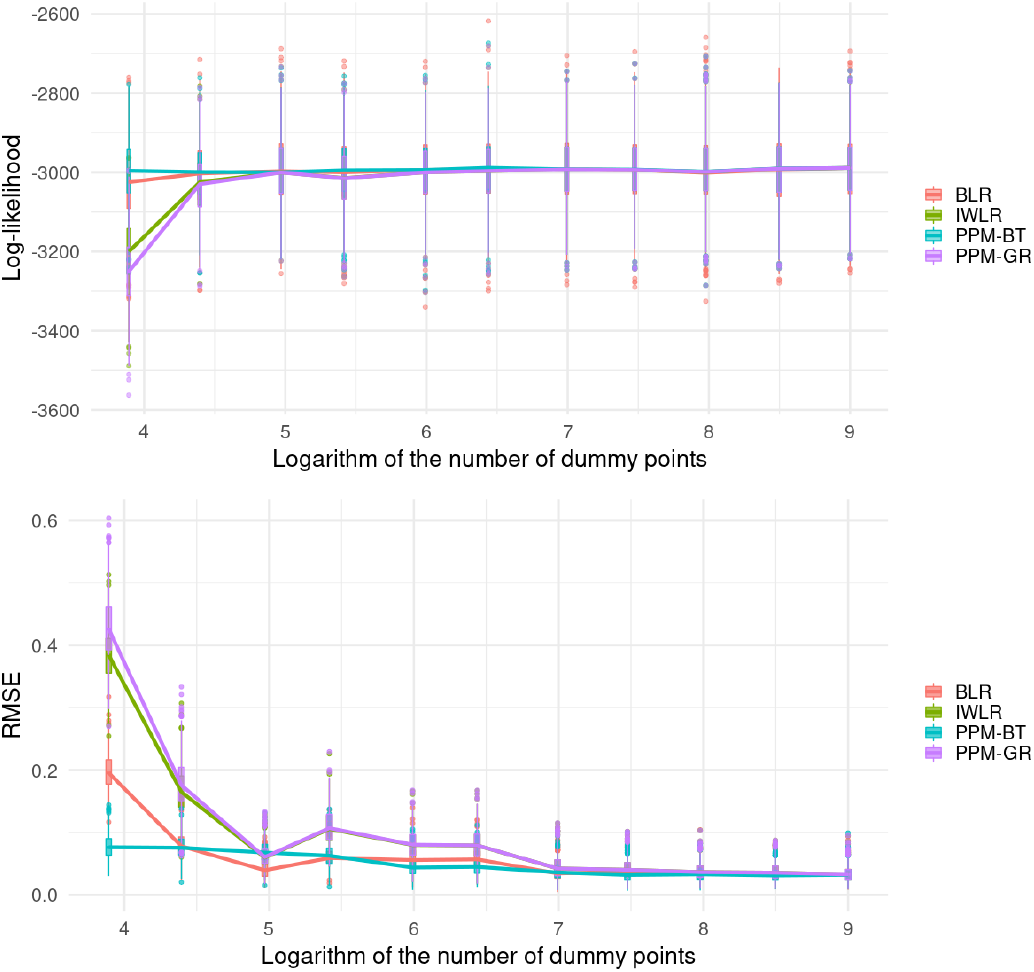
Log-likelihood values and residual mean squared errors for various numbers of dummy points, using BLR, IWLR, PPM-GR and PPM-BT methods. Note that the log-likelihood for the BLR has been scaled and shifted.

## 5 Case Study

The case study aims to demonstrate how the Poisson point process framework is applied to a PO data set of *Arctotheca calendula* (commonly known as Capeweed) from 2010-2021 spanning Southern Australia (GBIF, 2021). This species was chosen as it does not appear to have been the subject of any SDM analysis in the current literature, and is relevant to Australian agriculture as it is a pest and competitor to legume crops (Conning et al., 2011) and leads to weight reduction for livestock in pastures where it is present (Brundu et al., 2015). The *Arctotheca calendula* data was modelled by a Poisson point process model driven by environmental factors known to affect its distribution. For variable and model selection the *Arctotheca calendula* PO data was partitioned into a a training set (6; 282 points) and a validation set (985 points). The training data set consisted in occurrence data collected between 2013-2021, while the validation data set was collected between 2010-2012. The study window was determined from the domain of the *Arctotheca calendula* training data, with a small buffer, in total spanning from 130° to 154° longitude and −44° to −30° latitude. Since *Arctotheca calendula* is a terrestrial species, the window was restricted to terrestrial areas.

In a similar manner to the simulation study, the model was fitted for various numbers of dummy points distributed on a rectangular grid. The number of dummy points varied between 20 and 440, 424. The two best methodologies, PPM-BT and BLR, were implemented by using the `ppm` function from the `spatstat` package (Baddeley et al., 2015). Whilst there are many possible fitting choices within this function, these were not utilised in this study in order to ensure a fair comparison between the methodologies. The AUC on the validation data was recorded for each model fit as a measure of model performance.

A selection of soil (Viscarra Rossel et al., 2014) and bioclimatic covariates (Fick and Hijmans, 2017) with biological relevance to *Arctotheca calendula* were initially chosen for the model, as there is evidence that including both soil and climatic variables can improve model performance over climatic variables only (Hageer et al., 2017). An additional precipitation covariate was also included to be considered for the final model, equal to the total precipitation over autumn. This covariate was constructed as none of the 19 bioclimatic variables sourced explicitly related to the autumn germination period for *Arctotheca calendula*. Since our dataset was opportunistically sampled, following Renner et al. (2015) we introduce an additional covariate account for observer sampling bias. We used accessibility to high-density urban centres (Weiss et al., 2018). Each covariate was normalised.

Likelihood ratio testing was employed to identify the most important bio-climatic covariates using the `anova.ppm` function in `spatstat` Baddeley et al. (2015). This was performed by constructing models with linear covariates, omitting a single covariate at a time and performing the likelihood ratio test in an iterative process. This led to the construction of the final model, with the inclusion of second order interaction terms between covariates. The variables finally selected and included in the Poisson PPM in addition to observer bias are BIO8, BIO14, BIO18, pH and the interaction between BIO8 and pH. The model fitting results are shown in Table 2.

**Table 1:** Bioclimatic and Soil Covariate Summary

| Covariate | Description |
| --- | --- |
| pH | Soil acidity |
| P | Phosphorus concentration |
| Sand | Sand concentration |
| BIO6 | Minimum temperature of warmest month |
| BIO8 | Mean temperature of wettest quarter |
| BIO9 | Mean temperature of driest quarter |
| BIO10 | Mean temperature of warmest quarter |
| BIO11 | Mean temperature of coldest quarter |
| BIO13 | Precipitation of wettest month |
| BIO14 | Precipitation of driest month |
| BIO16 | Precipitation of wettest quarter |
| BIO17 | Precipitation of driest quarter |
| BIO18 | Precipitation of warmest quarter |
| Autumn precipitation | Precipitation between March-May |
| Bias | Travel time to urban center |

**Table 2:** Parameter estimates for the final model, along with their 95% confidence intervals.

| Parameter | Estimate | CI95.lo | CI95.hi |
| --- | --- | --- | --- |
| Intercept | 1.28 | 1.17 | 1.38 |
| BIO8 | 0.07 | -0.04 | 0.17 |
| BIO14 | 0.38 | 0.25 | 0.50 |
| BIO18 | -1.44 | -1.61 | -1.28 |
| pH | -1.18 | -1.27 | -1.10 |
| Bias | -3.44 | -3.55 | -3.32 |
| BIO8:pH | -0.76 | -0.85 | -0.66 |

The results are essentially equivalent for the BLR and PPM-BT methodologies for a large amount of dummy points. The BLR method starts converging in Fig. 3 for the recommended minimum number of dummy points of 4 times the number of presences (reached for a number of dummy points of around exp(10)), providing support for the recommendation proposed in Baddeley et al. (2014) to be upheld. The AUC of the PPM-BT method does not change much with the number of background points, varying only by 0.25% at the maximum. In contrast, the AUC for the BLR method varies more, but varies only by 0.30% once the recommended number of dummy points is reached. The final AUC of both methods is quite similar and both are within 0.12% of one another, which is consistent with both methods having converged. As in the simulation study, the PPM-BT method significantly out-performs BLR when the number of dummy points is small (less than the presence points). The final model performance is shown in Fig. 4. The fitted intensity aligns fairly well with the presence only points of both the training and validation datasets.

**Figure 3:**
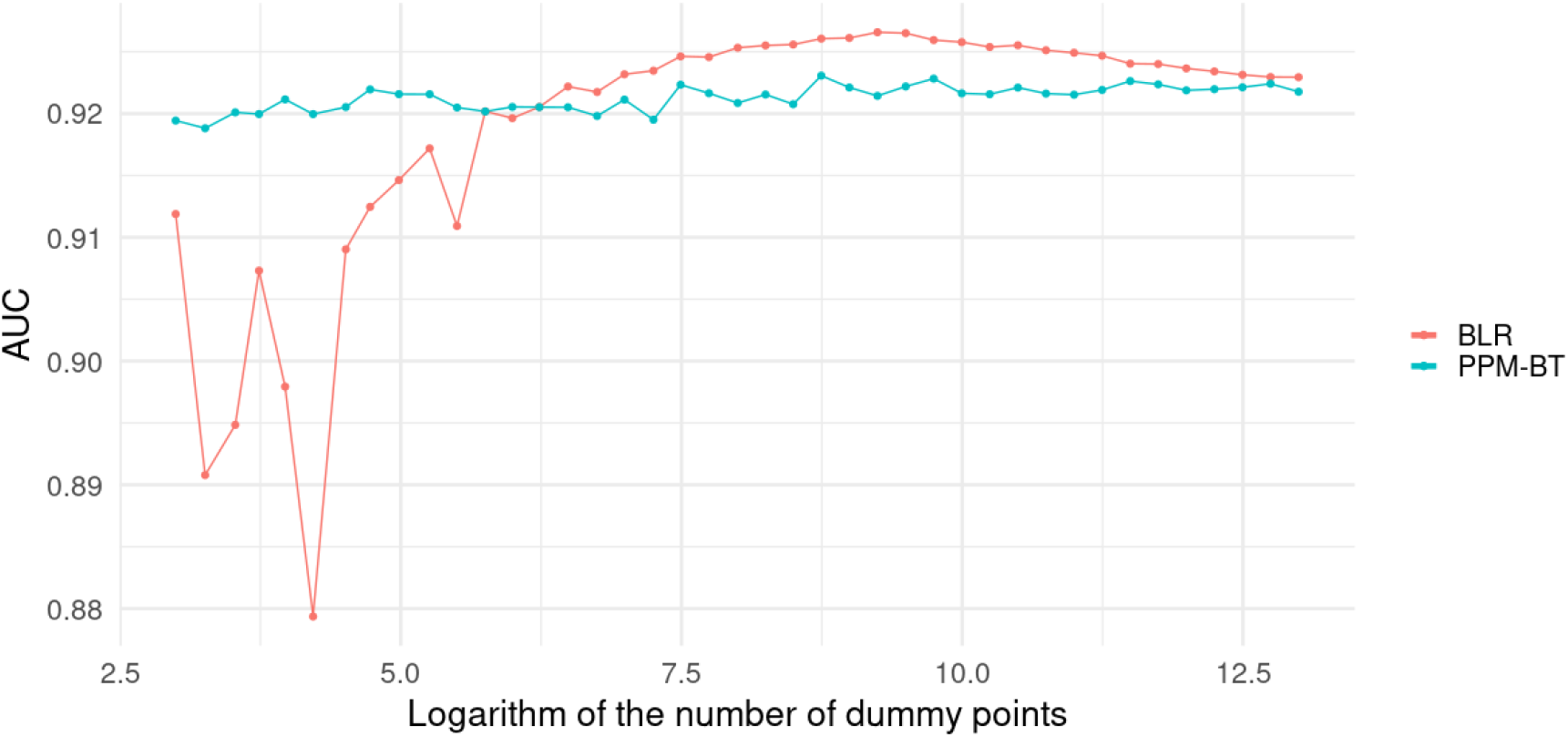
AUC against number of dummy points for the Berman-Turner device and Baddeley’s logistic regression methods.

**Figure 4:**
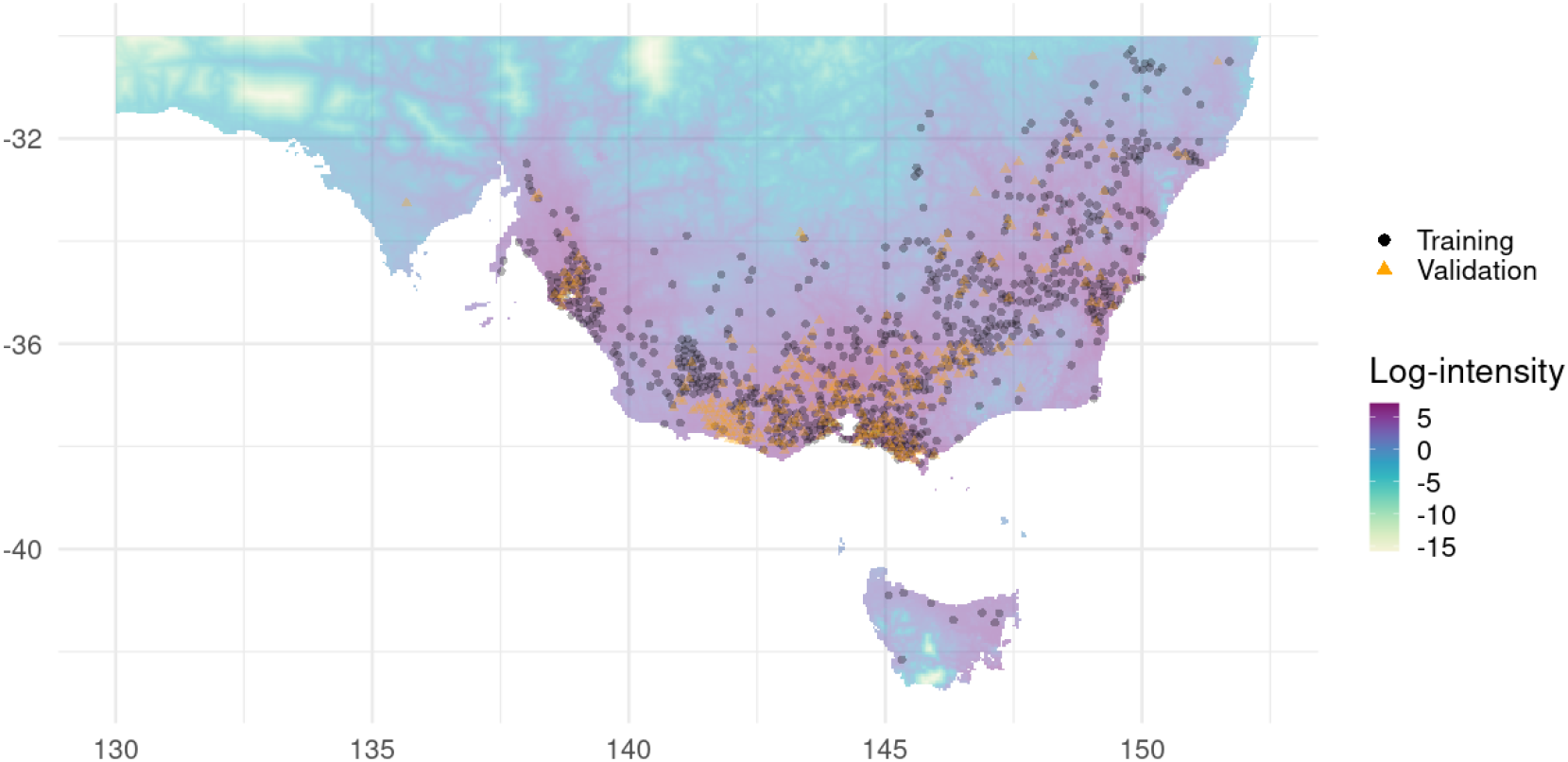
Fitted intensity of *Arctotheca calendula* based on the best fitted model using the PPM-BT and BLR methodologies, with training (black circles) and validation (orange triangles) data points shown. For clarity of presentation, the data was spatially filtered using the `ecospat` package (Broennimann et al., 2021) with a minimum nearest-neighbour distance set to 5 arcminutes.

## 6 Discussion

We have shown that the PPM-GR, PPM-BT, IWLR and BLR methods produce estimates that converge to the true parameter values as the number of dummy points increases (as the resolution of the grid upon which these points are located decreases). This means that all four methods are asymptotically equivalent and efficient in estimating the coefficient of the corresponding Poisson point model. However the accuracy of the estimates obtained from PPM-GR, IWLR and BLR rely significantly on the number of dummy points used. They converge to the true values at different rates and perform differently at coarse resolutions when there are too few dummy points. We did not find BLR to be the superior method. Rather, both the simulation and case study show that the PPM-BT outperforms BLR, especially under coarse resolutions. This observation is corroborated by our analysis of the RMSE, which is found to be significantly smaller for PPM-BT compared to the other methods at coarse resolutions. This therefore supports the notion that the Berman-Turner device is a more efficient option wherein a smaller number of back-ground/quadrature points is required to achieve satisfactory estimates. A potential explanation for the this method’s performance is that the Berman-Turner device also includes the presence points in addition to the dummy points in the numerical quadrature. All methodologies can be implemented with GLM software, therefore no advantages are conferred in this regard. Our results support the recommendation made by Baddeley et al. (2014) to generate four times as many dummy points as there are presence points, as clear convergence of the parameter values and RMSE at that point.

Contrary to the other three methods, Baddeley’s logistic regression (BLR) can also be used with general Gibbs point processes. Compared to Poisson point processes, points in a Gibbs point processes are correlated with one another, allowing them to model a wider range of phenomena, see e.g. Flint et al. (2022) for their application to ecology. Thus, although we found the performance of BLR less consistent than that of PPM-BT, BLR can be applied to a wider range of settings.

The similar behaviour of the IWLR and the PPM using gridded quadrature was not known before, and links this infinitely weighted logistic regression to the conventional Poisson PPM approach. This practical implications for researchers interested in implementing IWLR. Instead of fitting the IWLR’s log-likelilhood function in Eqn 1, one can simply fit the Poisson PPM method with the very simple quadrature scheme.

According to our knowledge, this is the first study to model the distribution of *Arctotheca calendula* through the PPM. There was obviously a significant observational bias in the *Arctotheca calendula* records obtained through GBIF datasets. The Poisson PPM with bias correction predicts the species’ distribution very well. Our modelling results show that Capeweed show superior durability to acidic soil, high temperature and low precipitation environments, which are consistent with ecological studies (Chapman et al., 2000). These factors, along with its greater moisture retention in seed coating, indicate that the germination period is the most important in respect to competition between *Arctotheca calendula* and *Trifolium subterraneum*, a domesticated annual pature legume as well as other domesticated annual pasture legumes (Conning et al., 2011). One limitation of a PPM is that there is residual short-range clustering that is not accounted for in a Poisson model. This is perhaps unsurprising, given reproductionF can occur through dispersal by wind and water, and vegetative propagation a potential mechanism for clustering (Brundu et al., 2015). An alternative approach is to fit the data with a Cox process (Renner et al., 2015), which is outside the scope of this study. Further research could investigate modelling choices to improve upon our presented model.

## 7 Conclusion

The study results support the implementation of the Berman-Turner device for Poisson point process modelling over other approaches using simple quadrature points, Baddeley’s logistic regression and the infinitely-weighted logistic regression methods. The Berman-Turner device produces more stable parameter estimates as the number of dummy points decreases, and superior estimates of the species intensity pointing towards a more robust fitting approach. The suggested minimum number of dummy points for BLR, which is four times the number of presence points, appears to be a sensible suggestion to achieve convergence of the parameter estimates. Users should be made aware though that this does not guarantee convergence of the estimates, and that increasing the number of dummy points beyond that threshold improves estimates.

